# MERFISH 2.0, an ultra-sensitive single-cell spatial transcriptomics imaging chemistry across diverse tissue types

**DOI:** 10.64898/2026.03.06.710199

**Authors:** Lizhi He, Bin Wang, Timothy Wiggin, Renchao Chen, Hao Wang, Bing Yang, Sudhir Gopal Tattikota, Lizi Maziashvili, Tong Zhang, Sainitya Revuru, Shichen Wang, Shradha Patil, Yijia Sun, Yi Sun, Man Li, Yuan Cai, Lingxian Wu, Natalia Pentrenko, Angela Vasaturo, Manisha Ray, George Emanuel, Jiang He

## Abstract

Spatial transcriptomics has emerged as a transformative approach for elucidating tissue architecture, cellular heterogeneity, and disease mechanisms by preserving the spatial context of gene expression in cells. Despite these advances, many spatial transcriptomic methods underperform in archival or clinically relevant specimens, particularly formalin-fixed, paraffin-embedded (FFPE) tissues, where RNA degradation and crosslinking hinder transcript detection. To address these challenges, we developed Multiplexed Error Robust Fluorescence In Situ Hybridization 2.0 (MERFISH 2.0), an optimized spatial transcriptomic imaging chemistry to enhance profiling of fragmented and highly crosslinked RNA. Across diverse human and mouse tissues preserved as fresh-frozen, fixed-frozen, and FFPE specimens, MERFISH 2.0 substantially increased transcript detection sensitivity by up to ∼8-fold relative to MERFISH 1.0 while preserving quantitative concordance (Pearson r ≥ 0.8 across tissues). In archived fresh-frozen human brain samples, MERFISH 2.0’s enhanced sensitivity improved transcript recovery, enhanced cell type resolution and spatial analyses. In low quality archival FFPE human breast cancer specimen, MERFISH 2.0 revealed additional cell populations, novel cell clusters, refined tumor–immune architecture, and increased detection of gene–gene and cell–cell interactions relative to MERFISH 1.0, underscoring the impact of improved sensitivity on downstream spatial analysis. By substantially expanding robust transcript detection to degraded and archival samples, MERFISH 2.0 enables scalable, cohort-level spatial transcriptomic analysis across clinically relevant tissue collections.

## Introduction

Over the past decade, dissociated single-cell and single-nucleus RNA sequencing technologies have fundamentally reshaped our understanding of cellular diversity and cell state in both normal and diseased tissues^1–5^. By resolving gene expression at cellular resolution, these approaches have enabled the discovery of new cell types, transient cell states, and lineage relationships^2,6–8^. However, the requirement for tissue dissociation disrupts native spatial architecture, eliminating information about cellular organization, microenvironmental context, and cell–cell interactions that are fundamental to understanding tissue function, disease progression, and drug treatment response^9–11^. Spatial transcriptomic approaches overcome this limitation by quantifying gene expression directly within intact tissue, thereby enabling direct interrogation of tissue architecture, spatially restricted gene programs, and intercellular communication *in situ*^12–15^.

Broadly, spatial transcriptomics methods fall into two categories: sequencing and imaging-based modalities^13,15–20^. Sequencing-based approaches provide genome-wide coverage and are valuable for unbiased discovery but are limited by per-transcript sensitivity and reduced robustness in degraded material^21^. Furthermore, sequencing-based approaches, in most implementations, lack true single-cell resolution as the captured spot size does not match cellular dimensions^22^.

Imaging-based spatial transcriptomic technologies instead are targeted, and interrogate a defined set of biologically and disease-relevant transcripts, while achieving true single cell resolution^15,18,23,24^, enhanced sensitivity^18,22,24,25^, reproducibility^26^, and quantitative accuracy^18,22,24,25^. This design is particularly well suited for hypothesis-driven studies, large cohort analysis, and translational applications where consistent measurement across samples, sites, and time points are required while still supporting discovery through identification of spatially defined programs and interactions.

Among imaging-based spatial transcriptomics technologies, multiplexed error-robust fluorescence in situ hybridization (MERFISH) is one of the most adopted technologies and widely implemented in different biological applications^15,20,22,24,27–29^. Beyond spatial transcriptomic imaging, MERFISH has been demonstrated to enable spatial epigenomic profiling^30^, isoform resolved whole-transcriptome imaging^31^, DNA chromatin tracing^32,33^, and three-dimensional imaging of thick tissues^34^. The Vizgen MERSCOPE and MERSCOPE Ultra platforms leverage MERFISH and provide an easy-to-use, integrated workflow to enable researchers to image up to 1000 genes with single cell and subcellular resolution for up to 3 cm^2^ sized specimen^35–37^. The platform also allows simultaneous detection of protein and RNA on the same tissue slices, enabling spatial multiomics measurements^38,39^. Researchers have leveraged the MERSCOPE platform to profile gene expression in cultured cells^40,41^, frozen and FFPE tissue samples, enabling the construction of cellular atlases of a variety of tissue types, including mouse brain^42–44^, mouse skin^45^, arabidopsis^46^, human oral tissue^47^, human skin^48^, and human brain^49,50^ among others. MERSCOPE assays have also been instrumental in identifying niches that predict immunotherapy response in hepatocellular carcinoma^51^, discovering spatially organized immune cell hubs in lung cancer linked to therapeutic outcome^52^, characterizing the mechanism of disease progression in AD^53,54^ and pinpointing which cells change, where they are located, and how immune–brain interactions drive pathology in encephalitis^55,56^.

Compared to other imaging-based spatial transcriptomic technologies, MERSCOPE has been cited to have the highest sensitivity and optimal trade-off between sensitivity and specificity when the RNA quality in the biological sample is high^57^. However, despite its demonstrated sensitivity and specificity in high-quality samples, RNA detection remains challenging in FFPE tissues^58^, which represent the dominant preservation method in clinical practice and a vast repository of archival specimens. Specifically, formalin-induced crosslinking reduces probe accessibility and increases background fluorescence, while RNA fragmentation limits probe binding sites and compromises transcript detection. These factors compromise detection sensitivity and downstream single-cell analyses in FFPE samples with low RNA quality, limiting the utility of original MERFISH chemistries (referred as MERFISH 1.0) in clinically relevant samples ^59,60^.

To address these constraints, we developed MERFISH 2.0, a novel spatial transcriptomic imaging chemistry and sample preparation workflow optimized for profiling fragmented and highly crosslinked RNA. Specifically, we first optimized RNA anchoring to increase retention of fragmented RNA transcripts during hydrogel embedding, so that more RNA targets are captured for downstream profiling. Second, we optimized the probe structure to enhance hybridization efficiency between RNA transcript target and detection probes. Thirdly, we introduced a controlled signal enhancement strategy to boost signal-to-noise ratio during imaging while preserving quantitative accuracy. Altogether, these changes allowed us to substantially improve transcript detection sensitivity relative to MERFISH 1.0 (up to ∼8-fold, depending on tissue and preservation method) while preserving quantitative concordance.

In this study, we benchmark MERFISH 2.0 against MERFISH 1.0 across a diverse set of human and mouse tissue types preserved as fresh-frozen, fixed-frozen or FFPE specimens. Using metrics including transcripts per unit tissue area (100 µm²), transcripts per cell, cross-chemistry concordance, and downstream biological performance in brain and tumor contexts, we demonstrate that enhanced sensitivity improves transcript recovery, reduces cell dropout, and strengthens single-cell spatial analyses. In low quality clinical samples, MERFISH 2.0 enables more comprehensive cell-type identification and reveals spatially organized cellular interactions that were previously obscured. By substantially improving sensitivity in degraded and crosslinked specimens while preserving quantitative fidelity, MERFISH 2.0 expands the applicability of spatial transcriptomics to translational and clinical research, unlocking access to the extensive FFPE tissue archives that underpin modern pathology.

## Results

### MERFISH 2.0 chemistry and workflow overview

MERFISH enables quantification of targeted RNA transcripts through combinatorial labeling, error-robust barcoding, and sequential imaging^15^. Each RNA transcript is hybridized with 30–50 encoding probes carrying a unique binary barcode that is decoded across imaging rounds. In MERFISH 1.0, transcripts are anchored to a hydrogel through their 3′ poly(A) tails^61^, and encoding probes tile along the RNA. The sample is then cleared using a matrix-imprinting-based clearing approach prior to imaging on the MERSCOPE^61^. For intact transcripts, this approach performs well because multiple probes binding along each RNA generate strong fluorescent signals during imaging (**Fig. 1A**). However, in lower-quality samples, including archival FFPE tissues or those with elevated RNase activity, RNA fragmentation and crosslinking frequently occur, leading to partial transcript loss after tissue clearing and reduced probe accessibility across the RNA sequence. The decrease in available binding sites may lead to diminished signal intensity, increased background fluorescence, and reduced accuracy in transcript detection and quantification (**Fig. 1B**).

**Figure 1.**
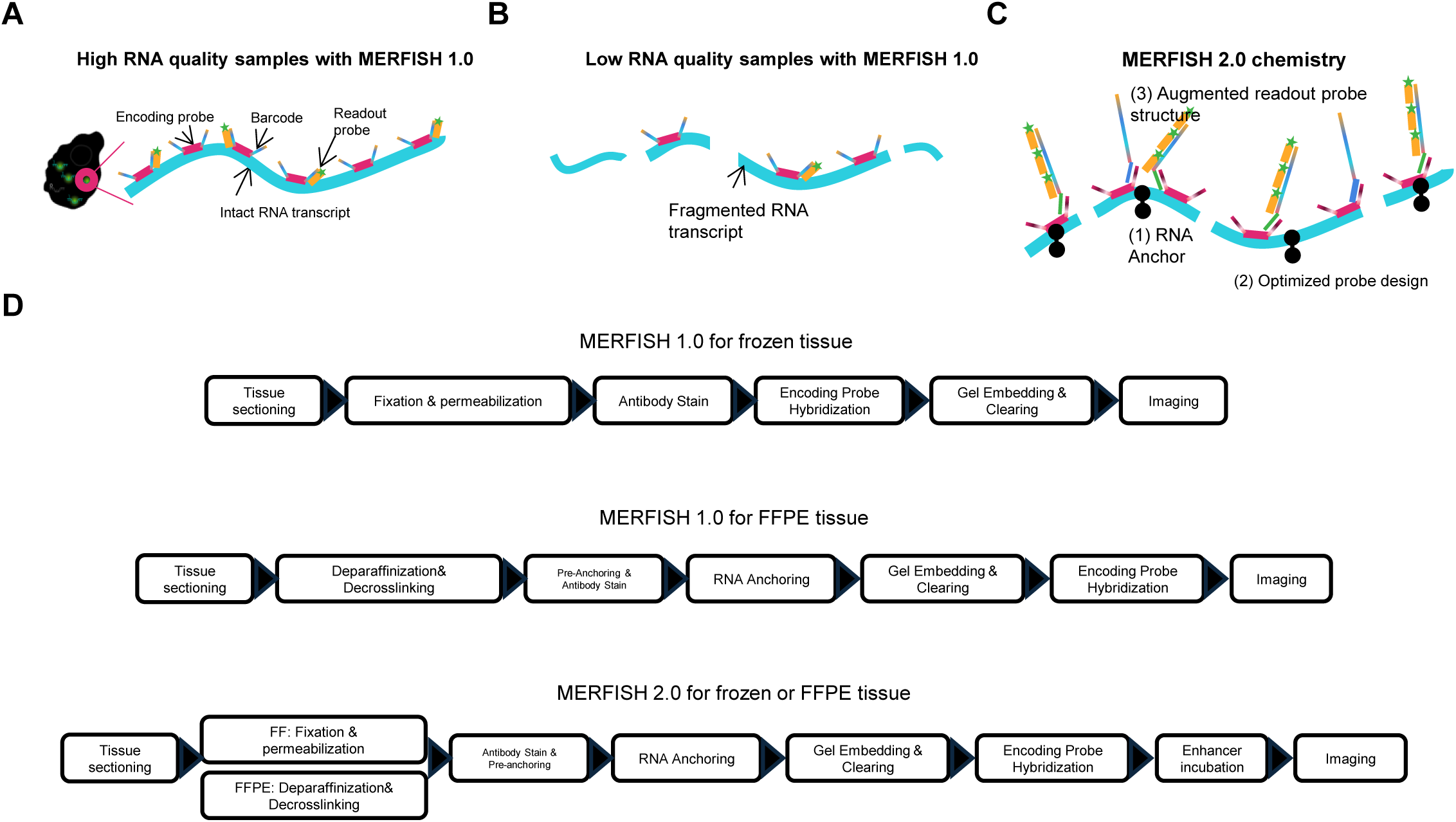
MERFISH 2.0 assay and workflow overview. **A**) Schematic of MERFISH 1.0 chemistry in samples with high RNA integrity. Intact RNA transcripts allow multiple encoding probes to bind along the transcript length, producing strong readout signals during imaging. **B**) Schematic of MERFISH 1.0 chemistry in samples with low RNA integrity. RNA fragmentation reduces the number of available probe-binding sites, leading to weaker fluorescence signals and decreased detection efficiency. **C**) Schematic of MERFISH 2.0 chemistry. MERFISH 2.0 workflow is developed to enhance RNA transcript detection efficiency in samples with low quality RNA. Improvements include optimized RNA anchoring to retain fragmented RNA transcripts during sample preparation, an improved encoding probe design for efficient hybridization, and augmented readout probe structure with an enhancer to increase imaging signal-to-noise ratio. **D**) Comparison of MERFISH workflows. Top, MERFISH 1.0 workflow for frozen tissue. Middle, MERFISH 1.0 workflow adapted for FFPE tissue. Bottom, MERFISH 2.0 workflow applicable to both frozen and FFPE samples.

To overcome these limitations, MERFISH 2.0 chemistry was developed to enhance transcript detection in samples characterized by RNA fragmentation and/or crosslinking. Key improvements include: (1) optimized RNA anchoring to more effectively immobilize fragmented transcripts within the hydrogel during sample preparation, thereby retaining more RNA targets for profiling; (2) redesigned encoding probe structures to enable more efficient hybridization between RNA targets and detection probes; and (3) the introduction of enhancers to amplify imaging signals in a controlled manner and improve overall signal-to-noise performance (F**ig. 1C**).

Separately, two distinct workflows were used to process frozen and FFPE samples with MERFISH 1.0 protocol (**Fig. 1D**). For frozen samples, encoding probe hybridization was performed prior to gel embedding and tissue clearing, whereas FFPE samples were first gel-embedded and cleared, followed by encoding probe hybridization. Because fixation conditions vary across frozen samples and affect encoding probe hybridization efficiency, sample fixation and permeabilization is subject to optimization in MERFISH 1.0 protocol. In contrast, a cleared sample is generally expected to present fewer obstacles for probe binding. Therefore, in MERFISH 2.0, we adopted a workflow in which samples are first gel-embedded and cleared, followed by encoding probe hybridization (**Fig. 1D**). After histological steps, samples are labeled with cell boundary stain, RNA transcripts are then immobilized through anchoring reagents. The samples are gel embedded, cleared, and hybridized with encoding probes, followed by enhancers hybridization prior to imaging. Overall, this resulted in a unified sample preparation workflow for both frozen and FFPE samples.

### Benchmarking MERFISH 2.0 against MERFISH 1.0 in fresh frozen mouse brain samples

To evaluate the performance of MERFISH 2.0, we first benchmarked MERFISH 2.0 against MERFISH 1.0 using high-quality fresh frozen mouse brain—a sample type where MERFISH has been extensively applied and for which abundant reference data are available^20,42–44,62^. Across 3 mouse brain slices measured with the 815-plex MERSCOPE Pan Neuro Cell Type Panel, MERFISH 1.0 detected an average of ∼185 transcripts per 100 µm² tissue area, with a mean of 727 counts per cell, indicating successful measurement and high data quality. MERFISH 2.0 exhibited an approximately two-fold increase in sensitivity compared with MERFISH 1.0, with an average transcript counts per 100 µm² around 393. Notably, MERFISH 2.0 data had reduced variability across biological replicates (n=3), indicating improved sensitivity and consistency (**Fig. 2A**).

**Figure 2:**
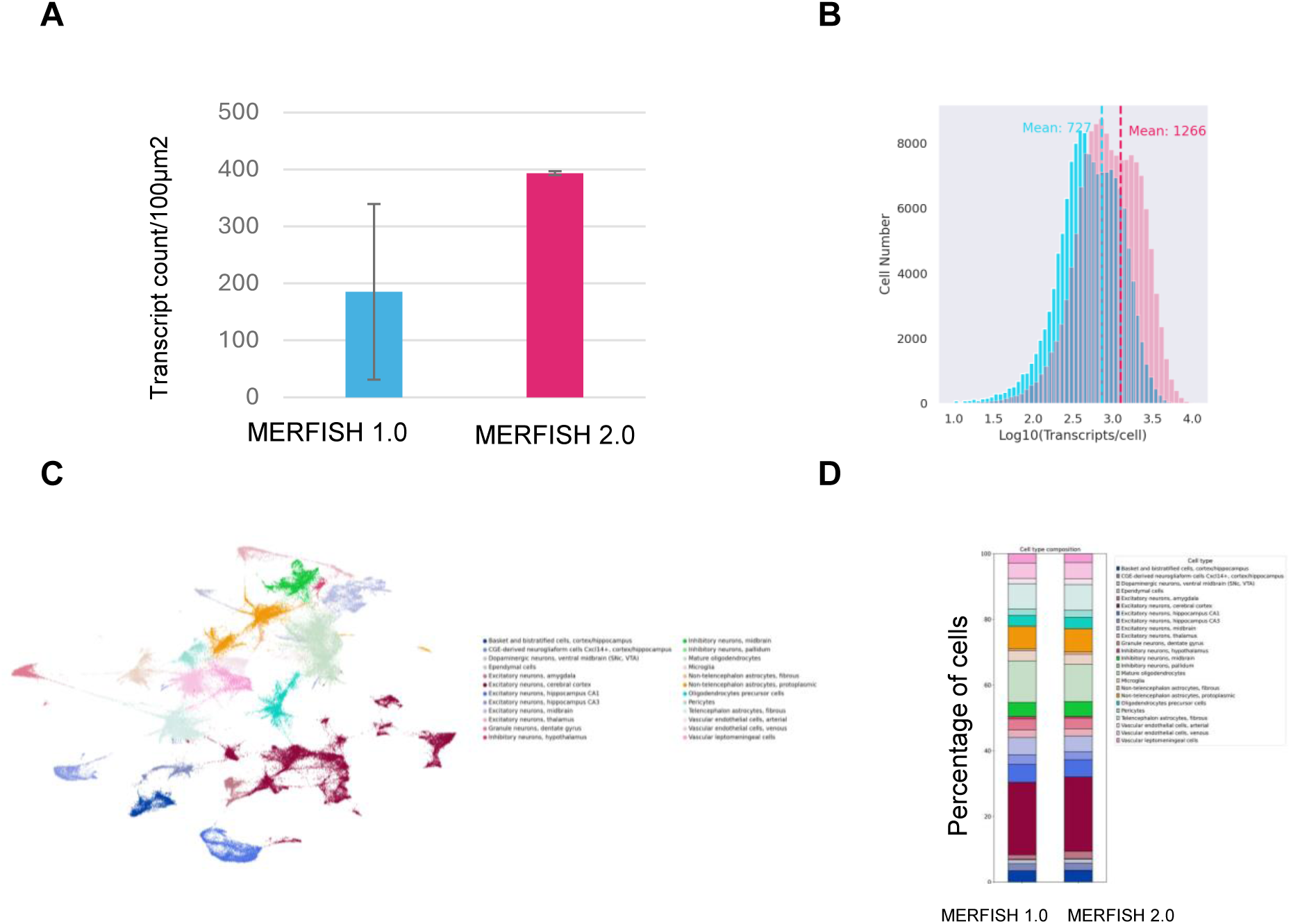
MERFISH 2.0 improves transcript detection and reduces variation in high quality fresh frozen mouse brain. Fresh frozen mouse brain sections were profiled with an 815-plex MERSCOPE Pan Neuro Panel and imaged on MERSCOPE platform to compare MERFISH 1.0 and MERFISH 2.0 chemistries. **A)** Sensitivity comparison between MERFISH (blue) and MERFISH 2.0 (red) measured as RNA transcript counts per 100 µm^2^.Bars represent mean values across biological replicates (N=3). MERFISH 2.0 shows increased transcript detection and reduced variability relative to MERFISH 1.0. **B**) Distribution of detected transcripts per cell for MERFISH 1.0 and MERFISH 2.0, shown as histograms of log10-transformed transcript counts per cell. MERFISH 2.0 exhibits higher transcript counts per cell. **C**) UMAP visualization of major neuronal and non-neuronal cell populations identified using MERFISH 1.0 and MERFISH 2.0 data. The MERFISH 1.0 and MERFISH 2.0 data was combined for analysis and co-embedding in the UMAP. D) Relative abundance of annotated cell types detected with MERFISH 1.0 and MERFISH 2.0, shown as the percentage of total cells assigned to each cell type.

At the single-cell level, MERFISH 2.0 yielded an increase of detected transcripts per cell, with a mean of 1266 counts per cell, reflecting higher molecular counts within cells (**Fig. 2B**).

MERFISH 1.0 and 2.0 data have excellent concordance, with correlation coefficient >0.9 (data not shown). This agreement is reflected in UMAP embeddings, where the same major cell types were identified in both datasets (**Fig. 2C**), and the relative proportions of major cell types were highly concordant between MERFISH 1.0 and MERFISH 2.0 (**Fig. 2D**). Furthermore, MERFISH 2.0 enabled improved cell subtype resolution in mouse brain relative to MERFISH 1.0, as the increased transcript counts per cell supported more refined subclustering analyses (data not shown). Because the primary goal of this study was to develop a chemistry compatible with lower-RNA-quality samples, we did not further investigate these improvements.

Together, these results demonstrate that MERFISH 2.0 enhances transcript detection efficiency, reduces run variations, without perturbing underlying cell-type composition in high-quality tissue.

### MERFISH 2.0 performance in fresh frozen human brain tissue

Because the human brain is also commonly used for benchmarking in spatial transcriptomics studies^62–64^, we extended our technical performance assessment to human brain tissue. Unlike mouse brain samples, human brain specimens are often collected postmortem, resulting in widely variable RNA quality. We used an 815-plex MERSCOPE Human Brain Panel to image the human brain.

Similar to the mouse brain, MERFISH 2.0 demonstrated substantially improved performance compared to MERFISH 1.0 in freshly collected human brain tissue, where RNA quality is anticipated to be high. Transcript density measurements showed a consistent increase in transcript count per tissue area, with MERFISH 2.0 detecting ∼2.5 fold more counts than MERFISH 1.0 (**Fig. 3A, 3E and 3F**). This increase produced a systematic upward shift in the scatter distribution of correlation analysis, with most genes lying above the x = y line (**Fig. 3A**). Measurements remained highly correlated across chemistry versions (r ≈ 0.91), indicating that MERFISH 2.0 provides a substantial gain in sensitivity while preserving quantitative consistency with MERFISH 1.0 (**Fig. 3A**). Furthermore, transcript counts from MERFISH 1.0 and 2.0 correlated well with bulk RNAseq data, with correlation coefficient at 0.7 for MERFISH 1.0 and 0.8 for MERFISH 2.0, respectively, suggesting accurate measurements (**Fig. 3C and 3D**).

**Figure 3:**
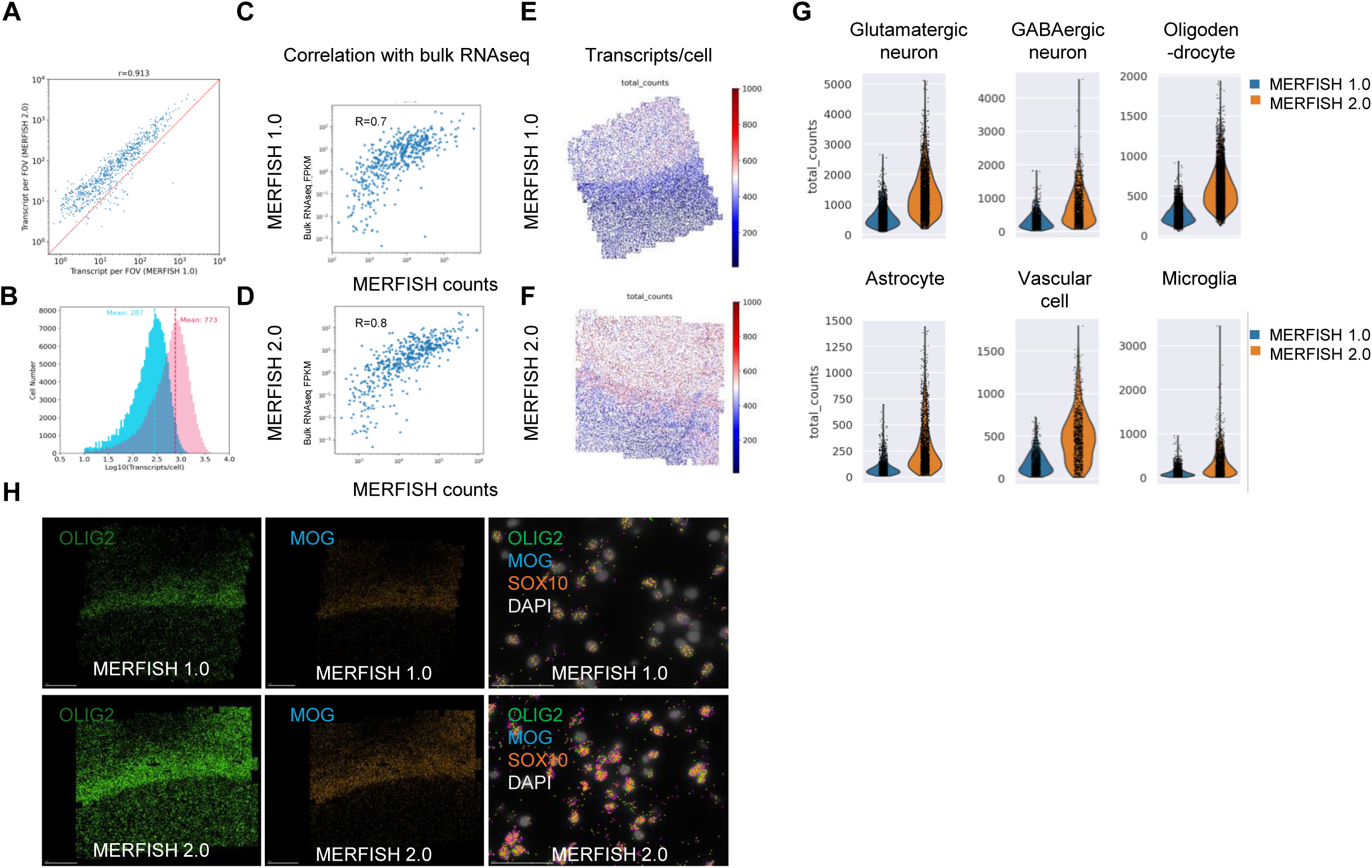
MERFISH 2.0 improves *in situ* gene expression profiling in high quality fresh frozen human brain samples. Fresh-frozen human brain sections were profiled using the 815-plex MERSCOPE Human Brain Panel and imaged on the MERSCOPE platform to compare MERFISH 1.0 and MERFISH 2.0 chemistries. **A)** Comparison of transcript detection between MERFISH 1.0 and MERFISH 2.0 on adjacent tissue slices. Scatter plot shows transcript counts detected by the two chemistries across the panel. **B)** Distribution of detected transcripts per cell for MERFISH 1.0 and MERFISH 2.0 shown as histograms of log10-transformed transcript counts per cell. MERFISH 2.0 shows increased transcript detection per cell. **C-D)** Correlation between bulk RNA-seq and MERFISH data with MERFISH 1.0 (C) or MERFISH 2.0 (D), demonstrating strong concordance between spatial and bulk measurements. **E-F)** Spatial heat map of transcript/cell across the tissue section for MERFISH 1.0 (E) and MERFISH 2.0 (F) chemistry illustrating increased transcript detection with MERFISH 2.0. **G)** Violin plot showing transcripts counts per cell across representative cell types identified in the human brain, including glutamatergic neurons, GABAegeric neurons, oligodendrocytes, astrocytes, vascular cells, and microglia cells. MERFISH 2.0 exhibits a broader dynamic range of transcript detection across cell types. **H)** Spatial distribution of representative marker genes with MERFISH 1.0 (top) and 2.0 (bottom) chemistry, including OLIG2, MOG and SOX10 with DAPI staining marking nuclei. MERFISH 2.0 shows substantially increased transcript counts for each marker gene.

At the single-cell level, MERFISH 2.0 yielded an approximately three-fold increase in transcript count per cell, with the mean count rising from ∼287 to ∼773 (**Fig. 3B**). This resulted in a substantial increase of dynamic range of transcript detection across all major cell populations, suggesting MERFISH 2.0 could enable the detection of lowly expressly genes in cells **(Fig. 3G).** To illustrate the sensitivity gain achieved with MERFISH 2.0, we visualized OLIG2, MOG and SOX10, three genes expressed in oligodendrocytes in the imaged samples at single cell resolution. There were substantially more detected transcripts per cell that overlapped with DAPI staining (**Fig. 3H**).

In archival fresh-frozen human brain tissue, where RNA quality is anticipated to be lower than freshly collected samples, MERFISH 2.0 produced larger gains: ∼5-fold higher transcript counts per 100 µm² and ∼2.5-fold higher mean transcripts per cell (95 vs 250 transcripts per cell for MERFISH 1.0 vs MERFISH 2.0, respectively; **Fig. 4A and 4B**). These improvements reduced cell dropouts and increased number of cells passing QC (10,463 vs 21,236 cells for MERFISH 1.0 vs MERFISH 2.0, respectively), enhancing spatial mapping of different cell types in the human brain (**Fig. 4C and 4D**). In addition, the dynamic range of gene expression across identified cell types was substantially broader in MERFISH 2.0 compared to MERFISH 1.0 (data not shown), which is expected to improve downstream single-cell analyses. Notably, improved transcript recovery resulted in a denser and more continuous spatial distribution of cell types, as fewer cells were excluded during preprocessing. Consequently, anatomical structures in the human brain were more clearly resolved, with cortical layers more distinctly delineated in the MERFISH 2.0 data (**Fig. 4C and 4D**). Consistent with this observation, MERFISH 2.0 enabled the identification of the MGE interneurons in the sample, which were not observed with MERFISH 1.0 (**Fig. 4E and 4F)**. Furthermore, among the major cell types identified in human brain, MERFISH 2.0 identified more astrocytes as compared to MERFISH 1.0 (**Fig. 4E and 4F**). Altogether, the data suggests that reduced cell dropout in MERFISH 2.0 enabled more accurate recovery of cell-type proportions, better reflecting the underlying biological composition of the human brain.

**Figure 4:**
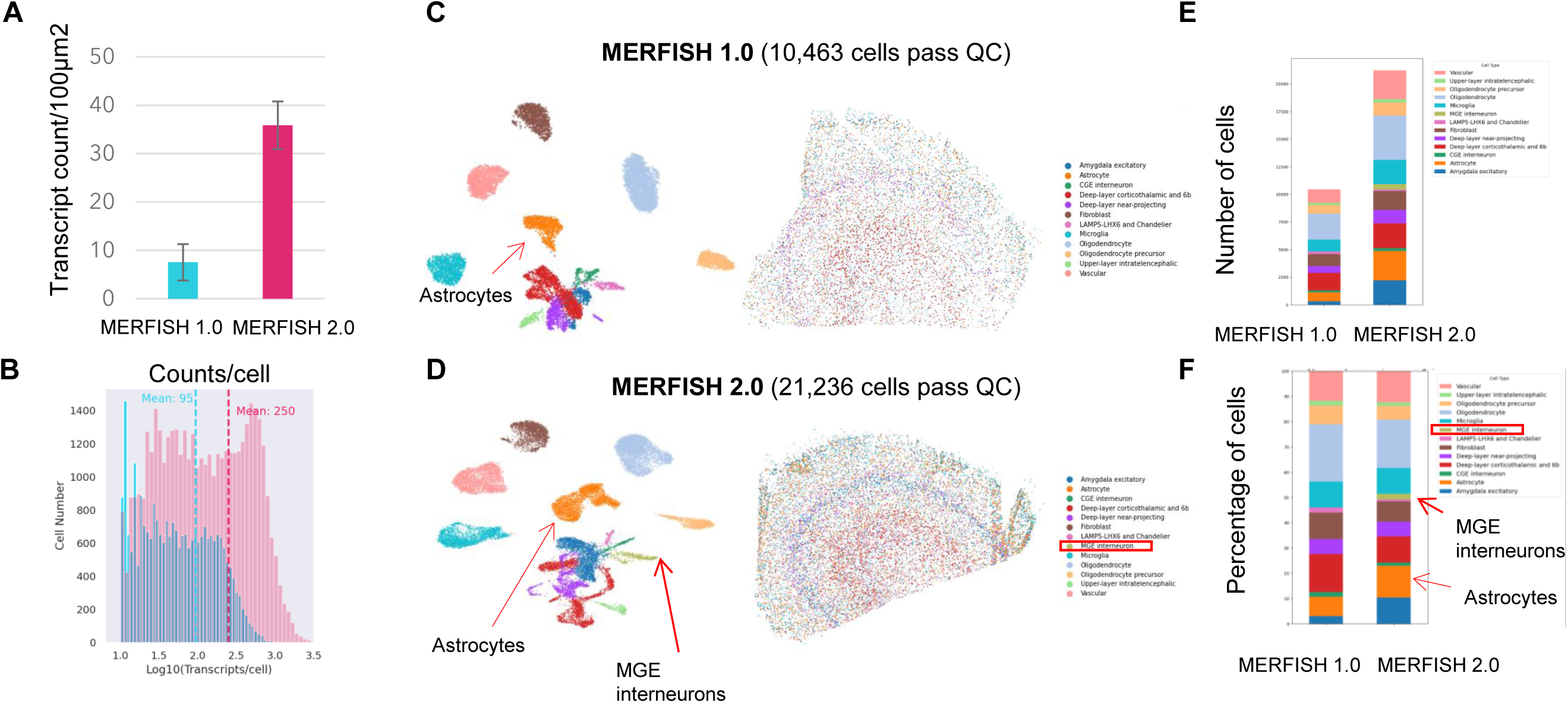
MERFISH 2.0 substantially improves transcript detection and cell recovery in low quality fresh frozen human brain samples. Archival fresh-frozen human brain sections with reduced RNA quality were profiled using the 815-plex MERSCOPE Human Brain Panel and imaged on the MERSCOPE platform to compare MERFISH 1.0 and MERFISH 2.0 chemistries. **A)** Sensitivity comparison between MERFISH 1.0 (blue) or MERFISH 2.0 (red), measured as RNA transcript counts per 100 µm². Bars represent mean values across biological replicates (n = 3). MERFISH 2.0 shows substantially increased transcript detection. **B)** Distribution of detected transcripts per cell for MERFISH 1.0 and MERFISH 2.0, shown as histograms of log10-transformed transcript counts per cell, demonstrating increased transcript detection with MERFISH 2.0. **C)** UMAP visualization of different cell types identified in human brain with MERFISH 1.0, with 10,463 cells pass QC. Left: clustering of cell types based on gene expression profiles. Right: spatial distribution of annotated cell types across the tissue section. **D)** UMAP visualization of cell populations identified in human brain with MERFISH 2.0, with 21,236 cells pass QC. Left: clustering of cell types. Right: spatial distribution of annotated cell types. MERFISH 2.0 recovers substantially more cells and improves detection of distinct cell populations. **E)** Absolute number of cells assigned to each cell-type cluster with MERFISH 1.0 and MERFISH 2.0 chemistry. **F)** Relative abundance of cell types detected with MERFISH 1.0 and MERFISH 2.0 chemistry, shown as the percentage of total cells assigned to each cluster. MERFISH 2.0 identifies a higher fraction of astrocytes and uniquely detects MGE interneurons that are not recovered with MERFISH 1.0.

### MERFISH 2.0 improves *in situ* gene expression profiling in multiple fixed frozen mouse tissue types

Having established the performance of MERFISH 2.0 in fresh frozen samples, we next evaluated the performance of MERFISH 2.0 in fixed frozen samples. We used an 815-plex Pan Mouse Panel and profiled a variety of fixed frozen mouse samples, including mouse liver, heart, kidney, lung, small intestine, brain and spleen. Consistent to what was observed in fresh frozen samples, transcript density was consistently higher with MERFISH 2.0 compared to MERFISH 1.0 across all tissues examined (**Fig. 5A**). The magnitude of improvement varied across tissues, however, with overall sensitivity increased by approximately two to three-fold. In addition, we observed excellent concordance between technical replicates for MERFISH 2.0, with correlation coefficients approaching 1 for all tissues (**Fig. 5B**). MERFISH 2.0 data also showed strong correlation with MERFISH 1.0 in these fixed frozen mouse tissues, with correlation coefficients exceeding 0.8 (**Fig. 5C**).

**Figure 5:**
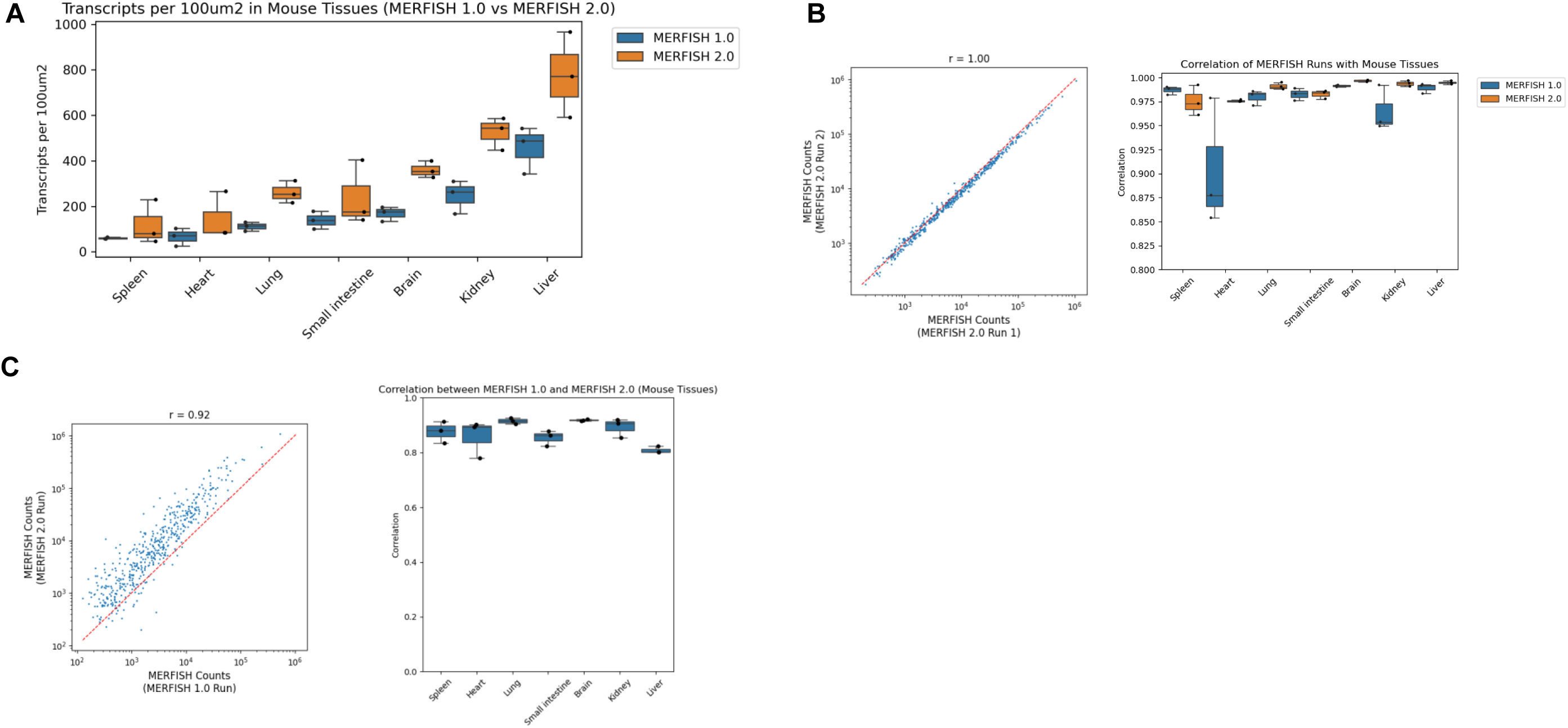
MERFISH 2.0 improves *in situ* gene expression profiling in multiple fixed frozen mouse tissue types. Fixed-frozen mouse tissues (spleen, heart, lung, small intestine, brain, kidney, liver) were profiled using the 815-plex Pan Mouse Panel following MERFISH 1.0 and workflows to evaluate assay performance across multiple tissue types. **A)** Sensitivity comparison between MERFISH 1.0 and 2.0. RNA transcript counts per 100 µm² were used as a metric of sensitivity. Each condition includes at least 3 biological replicates (N=3). MERFISH 2.0 consistently increases transcript detection across tissues. **B)** Reproducibility analysis of MERFISH 2.0 measurements across technical replicates. Left: representative scatter plot comparing transcript counts between two technical replicate MERFISH 2.0 runs. Right: Pearson correlation coefficients for replicate measurements across mouse tissues, demonstrating high reproducibility of MERFISH 2.0. **C)** Correlation between MERFISH 1.0 and MERFISH 2.0 measurements across tissues. Left: representative scatter plot comparing transcript counts detected by MERFISH 1.0 and MERFISH 2.0. Right: Pearson correlation coefficients calculated for each tissue type, showing strong concordance (r > 0.8) between the two chemistries.

Among the fixed frozen tissues profiled, mouse liver exhibited the highest transcript density per tissue area, with transcript counts per 100 µm² approximately 800 in the measured slices using this gene panel, whereas mouse spleen showed the lowest transcript density (**Fig. 5A**). This difference likely reflects the gene composition of the panel, which included fewer genes associated with spleen-specific cell types. Overall, these results establish robust performance improvements of MERFISH 2.0 in fixed frozen tissues and motivate subsequent evaluation in heavily fixed human samples.

### MERFISH 2.0 substantially improves sensitivity and single cell analysis in archival FFPE specimens

Next, we focused on evaluating whether MERFISH 2.0 improves sensitivity in archival FFPE human samples that exhibited low transcript counts when processed with the MERFISH 1.0 chemistry. FFPE samples present additional challenges for RNA imaging due to RNA fragmentation and chemical modification introduced during fixation and embedding, which can substantially reduce transcript recovery. To evaluate the performance of MERFISH 2.0 in FFPE samples, we created tissue microarrays (TMA) encompassing multiple cores of human breast cancer and lung cancer tissue from different patients. This allowed us to more efficiently evaluate the performance of MERFISH 2.0 in the same sample type with different RNA quality. Transcript counts per 100 µm^2^ was compared between MERFISH 1.0 and MERFISH 2.0 across multiple human tissue types, as well as at the individual core level. Similarly to our observations in frozen samples, MERFISH 2.0 improved the sensitivity for transcript detection across all tested samples (**Fig. 6A**). The largest gains were observed in low-quality TMA cores, where transcript densities increased up to 8-fold relative to MERFISH 1.0 (**Fig. 6A**). Correlation between MERFISH 2.0 and 1.0 data at the transcript level was high, with correlation coefficients above 0.83 for all profiled samples (**Fig. 6B**). Overall, this data suggests that MERFISH 2.0 was able to substantially improve sensitivity for RNA detection in FFPE samples.

**Figure 6:**
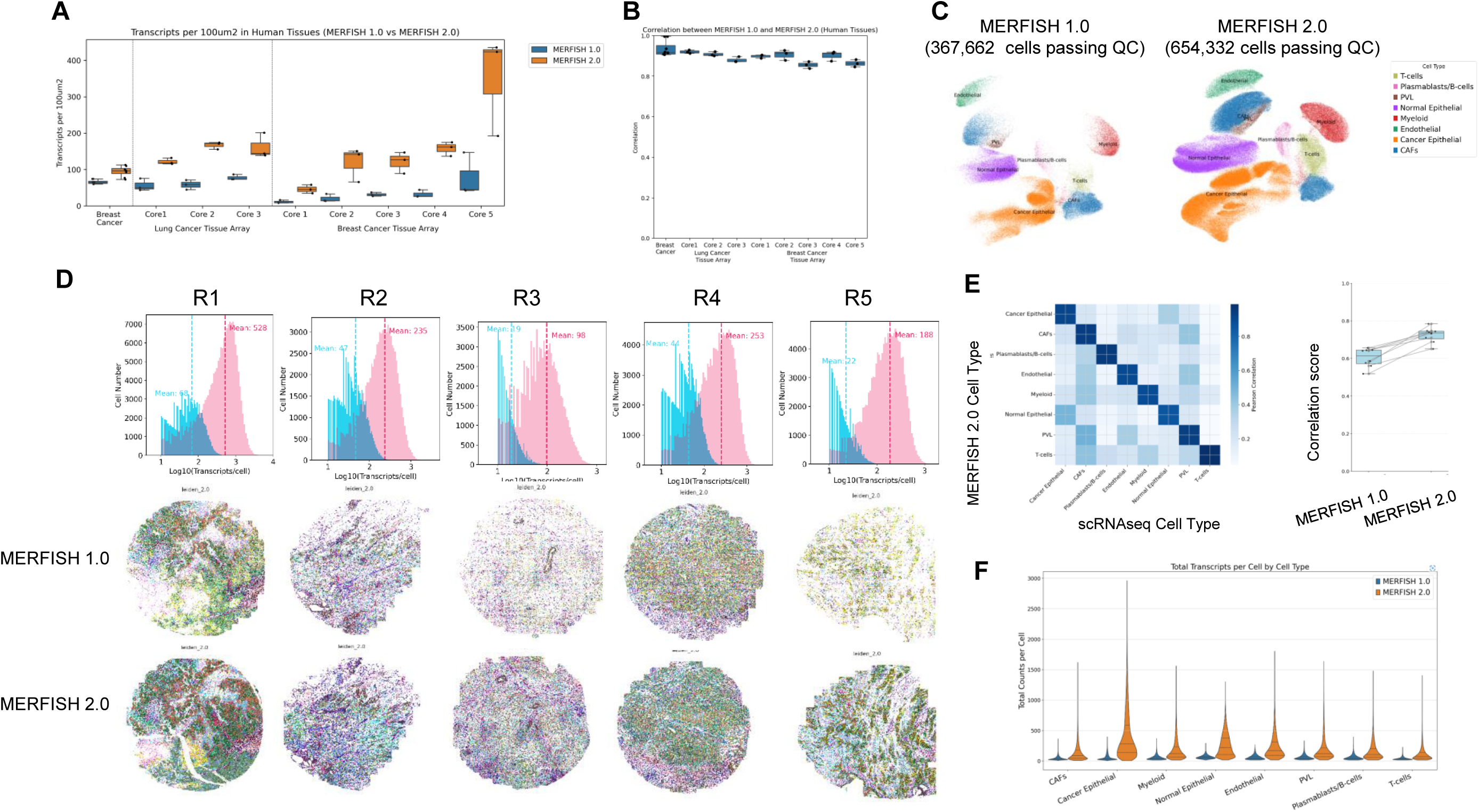
MERFISH 2.0 substantially improves sensitivity and single cell analysis in archival FFPE specimens. Formalin-fixed paraffin-embedded (FFPE) human tissues, including breast cancer specimens and lung and breast cancer tissue microarrays (TMAs), were profiled using the 815-plex Human Breast Cancer Panel or the Human Immuno-Oncology Panel following the MERFISH 1.0 and MERFISH 2.0 workflows. **A)** Sensitivity comparison between MERFISH 1.0 and 2.0 across FFPE human samples. RNA transcript counts per 100 µm² were used as a metric of sensitivity. Each condition includes at least 3 biological replicates (N=3). MERFISH 2.0 consistently increases the sensitivity for transcripts detection across samples. **B)** Concordance analysis between MERFISH 1.0 and MERFISH 2.0 measurements within each tissue. Person correlation coefficients calculated across samples demonstrate strong agreement between the two chemistries (r > 0.8). **C)**. UMAP visualization of single-cell transcriptomic profiles from FFPE human breast cancer displaying the different major cell types identified with MERFISH 1.0 (left), and MERFISH 2.0 (right). A total of 367,662 cells passed QC for MERFISH 1.0 and 654,332 cells passed QC for MERFISH 2.0. **D)** Transcript detection across individual TMA cores. Top, histograms showing the distribution of transcripts per cell for MERFISH 1.0 and MERFISH 2.0 across cores (R1–R5), with mean transcript counts indicated. Bottom, spatial distribution of Leiden clusters across tissue cores detected with MERFISH 1.0 (middle) and 2.0 chemistry (bottom), demonstrating much greater transcript coverage and cell recovery with MERFISH 2.0. **E)** Concordance between MERFISH-derived cell type profiles and reference single cell RNA sequencing data. Heatmap and summary correlation analysis show improved agreement with scRNA-seq when using MERFISH 2.0 compared to MERFISH 1.0. **F)** Violin plot showing the dynamic range of RNA transcripts counts per cell across representative cell types identified in human breast cancer.

To determine whether the improved sensitivity yielded additional biological insights, we analyzed the archival human breast cancer TMA cores. Across the 5 different cores (R1-R5), the average number of detected transcripts per cell ranged from 22 to 68 with MERFISH 1.0, whereas MERFISH 2.0 achieved a 5-8 fold increase in transcript detection (**Fig. 6D**). The increased transcript recovery also enabled more cells to passe QC with 654,332 cells retained in MERFISH 2.0 dataset compared to 367,662 in MERFISH 1.0 dataset (**Fig. 6C**). This resulted in a substantially denser spatial distribution of cell types in each core with MERFISH 2.0 (**Fig. 6D**).

At the single cell level, there was high concordance between MERFISH 2.0 and 1.0 data for all major cell types identified (**Fig. 6E**). However, among the major cell classes, we identified many more T cells in MERFISH 2.0 data (**Fig. 6C**). MERFISH 2.0 data also correlated better with the scRNAseq data than MERFISH 1.0 data (**Fig. 6E**), due to its improved sensitivity. Furthermore, the dynamic range of gene expression for all cell types identified in human breast cancer was much broader with MERFISH 2.0, suggesting MERFISH 2.0 is more capable of detecting lowly expressed genes or rare transcripts in tissue than MERFISH 1.0 (**Fig. 6F**).

Overall, the data suggested that MERFISH 2.0 enhances sensitivity without compromising quantitative fidelity, even in challenging FFPE samples and enables more accurate biological investigation.

### MERFISH 2.0 substantially improves immunoprofiling and cell-cell interaction analysis in archival FFPE specimens

Tumor heterogeneity is a fundamental feature of cancer and is particularly well characterized in breast cancer, which comprises multiple molecular subtypes, including Luminal A, Luminal B, HER2-positive, and triple-negative breast cancer, that differ in clinical behavior, prognosis, and therapeutic response^65^. The presence of these well-defined subtypes, together with the extensive availability of archived FFPE clinical specimens, makes breast cancer a practical and informative model for evaluating spatial transcriptomic technologies and investigating tumor heterogeneity in situ^66,67^. Within this context, we further evaluated the performance of MERFISH 2.0 in immunoprofiling of the tumor microenvironment in the FFPE human breast cancer samples.

With increased transcript counts per cell, a larger fraction of cells passed quality control in the MERFISH 2.0 dataset, resulting in a more densely resolved cellular landscape in the human breast cancer sample (**Fig. 7A**). Notably, a greater number of T cells were detected in the stromal region of the selected tumor core (**Fig. 7A**). Correspondingly, the expression of canonical T-cell marker genes, including CD8A, CD4, and FOXP3, was markedly higher in the MERFISH 2.0 dataset compared with MERFISH 1.0 (**Fig. 7A, B**). To further illustrate the gain in sensitivity for these markers, we annotated major cell types based on gene expression profiles and visualized the spatial distribution of representative transcripts within a single field of view. As shown in **Fig. 7B**, MERFISH 1.0 detected few or no transcripts for several immune cell markers— including those for T cells, B cells, plasmablasts, and myeloid cells—whereas substantially higher transcript counts were observed with MERFISH 2.0. This improvement is particularly important for immunoprofiling of individual cells within the tumor microenvironment, where many marker genes are expressed at relatively low levels and improved detection sensitivity enables more accurate identification of immune cell populations.

**Figure 7:**
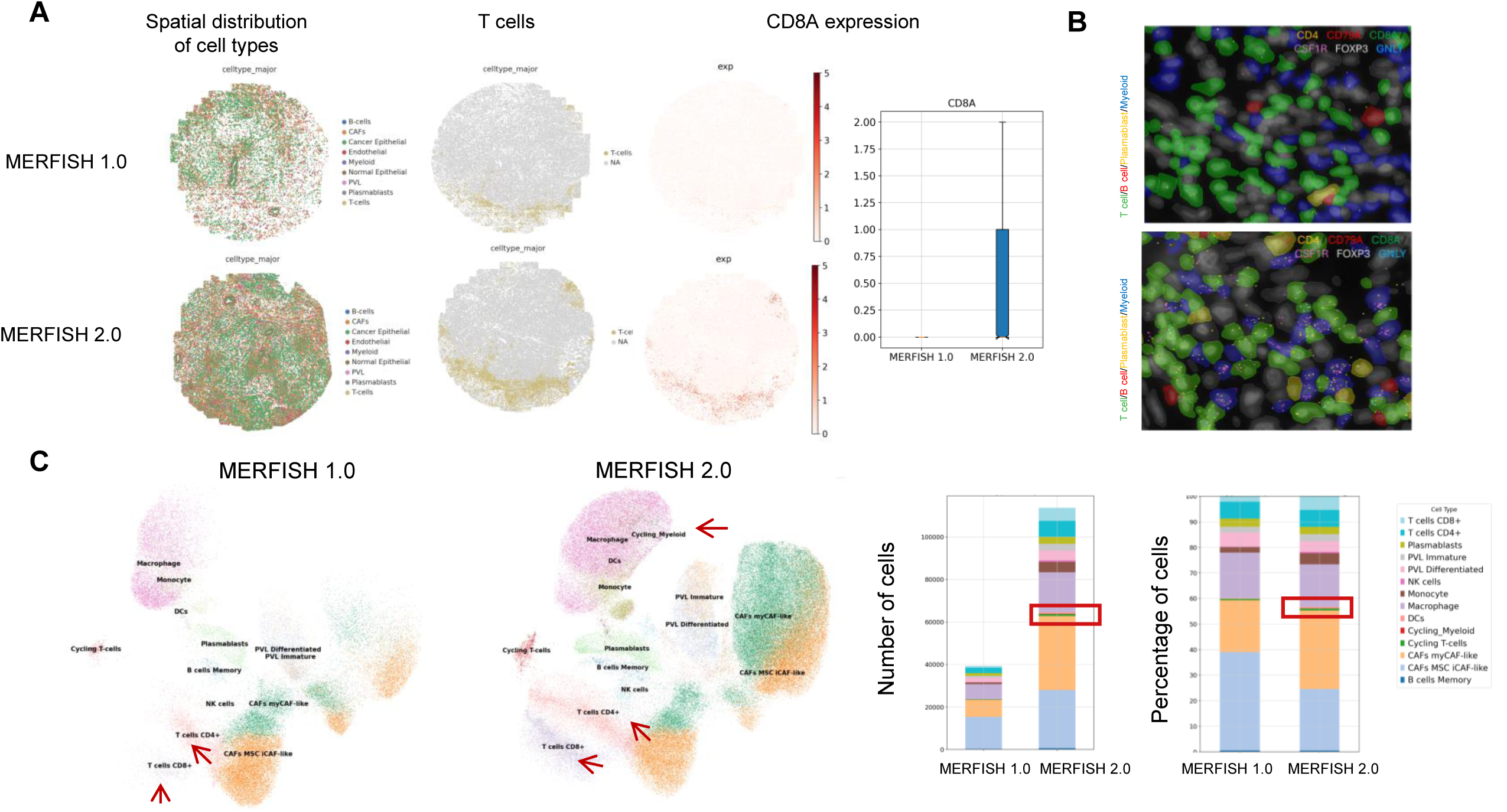
MERFISH 2.0 improves immunoprofiling of tumor microenvironment in FFPE human breast cancer. Immune cell populations from an FFPE human breast cancer sample were analyzed to evaluate whether the increased sensitivity of MERFISH 2.0 enables improved identification of immune cell subtypes. **A)** Spatial mapping of major cell populations and T cells. Left: spatial distribution of major cell types in the human breast cancer tissue profiled with MERFISH 1.0 (top) and 2.0 (bottom); Middle: spatial distribution of T cells identified in each dataset. Right: spatial distribution of CD8A gene expression and corresponding quantification of transcript counts showing increased detection with MERFISH 2.0 chemistry; **B**) Representative multiplexed images showing spatial localization of selected immune markers detected with MERFISH 1.0 (top) and MERFISH 2.0 (bottom) in annotated T cells (green), B cells (red), plasmablast cells (yellow) and myeloid cells (blue). MERFISH 2.0 detects substantially higher transcript counts for select marker genes across immune cell populations. **C**) UMAP visualization of immune cell subtypes identified with MERFISH 1.0 (left) and MERFISH 2.0 (right). Bar charts show the absolute number and proportion of cells assigned to each immune cell cluster. MERFISH 2.0 identifies larger populations of CD4⁺ and CD8⁺ T cells and reveals additional immune subtypes, including cycling myeloid cells that are not detected with MERFISH 1.0.

Next, we performed subclustering analysis to evaluate whether the improved sensitivity of MERFISH 2.0 could enable better resolution of cell subtypes within the tumor microenvironment. Immune and immune-associated cells were grouped from the dataset and subjected to further analysis. Consistent with the increased detection sensitivity, MERFISH 2.0 captured a larger number of CD4⁺ and CD8⁺ T cells compared with MERFISH 1.0. In addition, MERFISH 2.0 identified a distinct population of cycling myeloid cells that was not detected in the MERFISH 1.0 dataset (**Fig. 7C**).

With more cells retained after quality control, we anticipated that MERFISH 2.0 would improve the characterization of cell–cell interactions within the tumor microenvironment. To test this, we performed neighborhood analysis of major cell types following cell-type annotation to determine whether specific cell types are preferentially closer to each other in space. In the MERFISH 1.0 dataset, intercellular interaction patterns were much less pronounced, largely due to the smaller number of cells captured (**Fig. 8A**). In contrast, MERFISH 2.0 revealed clear spatial enrichment patterns, including close associations between B cells and T cells, as well as between endothelial cells and perivascular-like (PVL) cells (**Fig. 8A, B**). Together, these results suggest that the enhanced sensitivity of MERFISH 2.0 improves spatial analysis of cellular interactions within the tumor microenvironment.

**Figure 8:**
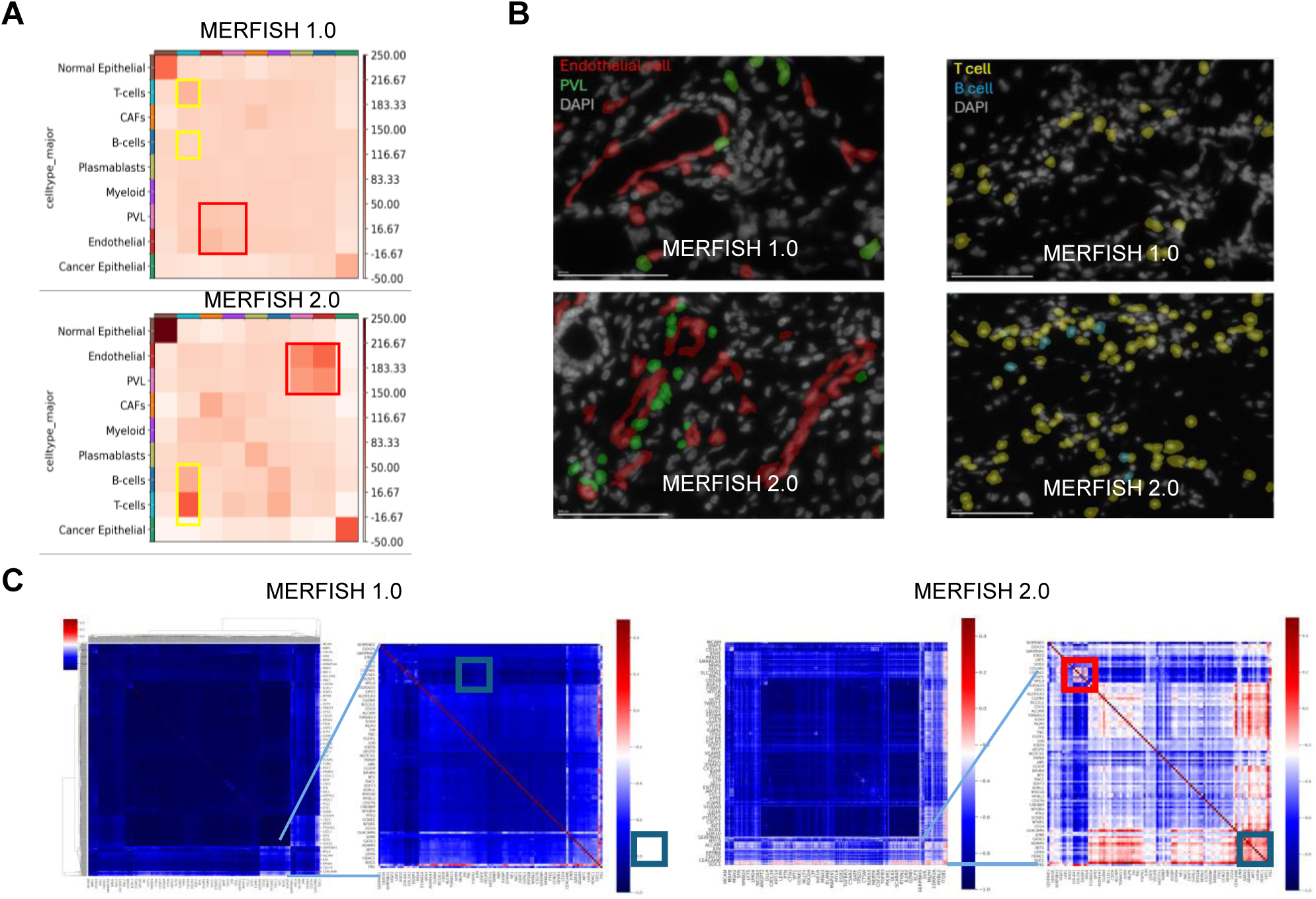
MERFISH 2.0 improves spatial interaction and gene-gene correlation analyses in FFPE human breast cancer. Spatial analyses were performed on FFPE human breast cancer samples to evaluate whether increased transcript detection with MERFISH 2.0 improves the characterization of cellular interactions and gene co-expression patterns. **A)** Spatial enrichment analysis of the major cell types identified with MERFISH 1.0 (top) and MERFISH 2.0 (bottom). Heatmaps show the degree of spatial association between cell types. MERFISH 2.0 reveals greater and more diverse spatial interactions among cell populations. **B**) Representative images illustrating spatial relationships between selected cell types. Left: interaction between endothelial cells and perivascular-like (PVL) cells; right: interaction between T cells and B cells only visible with MERFISH 2.0 (bottom) as B-cells did not appear with MERFISH 1.0 profiling. MERFISH 2.0 reveals these spatial relationships more clearly through enhanced transcript detection and improved cell-type annotations. **C**) Gene-gene correlation matrices from MERFISH 1.0 (left) and MERFISH 2.0 (right) datasets. MERFISH 2.0 reveals a larger number of correlated gene modules, indicating improved detection of coordinated gene expression programs.

Finally, gene-gene correlation analysis was performed to identify co-regulated gene modules in the human breast cancer samples. Owing to its higher detection efficiency, MERFISH 2.0 strengthened gene-gene correlation signals and revealed additional co-regulated modules compared with MERFISH 1.0 (**Fig. 8C**). These included genes involved in key signaling pathways, such as STAT2, JUN, and STAT6. Collagen genes also showed coordinated correlations in the MERFISH 2.0 dataset, consistent with extracellular matrix regulation in the tumor microenvironment. Notably, a correlation between GATA3 and AKT1 was detected, consistent with their known regulatory relationship in breast cancer. The increased recovery of gene correlations with MERFISH 2.0 enables improved inference of regulatory pathways and facilitates biomarker discovery and identification of potential therapeutic targets.

## Discussion

MERFISH is a quantitative and highly sensitive spatial transcriptomic imaging technology for *in situ* RNA profiling. Although it generates high-quality spatial transcriptomic data in well-preserved biological specimens, the original MERFISH chemistry is less optimized for low-quality samples and therefore exhibits reduced sensitivity under such conditions. This limitation restricts its application in clinical settings, where specimen quality is often variable.

Here, we demonstrate that MERFISH 2.0 substantially improves transcript detection sensitivity relative to MERFISH 1.0 across diverse preservation conditions, including fresh frozen, fixed frozen, and FFPE tissues, while maintaining high correlation in gene expression measurements between chemistry versions.

In high-quality frozen specimens such as mouse brain, MERFISH 2.0 increased transcript counts per area and maintained strong concordance with MERFISH 1.0 at both transcript and single-cell levels, indicating that sensitivity gains were achieved without compromising specificity. Notably, biological replicates exhibited reduced variability with MERFISH 2.0. This is likely to reflect improved detection efficiency, which reduces stochastic variation and enhances measurement consistency. Greater reproducibility across biological samples is advantageous for spatial data analysis, as it may mitigate batch effects in downstream analyses. Thus, even for high-quality samples, MERFISH 2.0 provides practical benefits over MERFISH 1.0.

In lower-quality specimens, such as postmortem human brain and FFPE samples, where RNA fragmentation and crosslinking reduce probe accessibility, MERFISH 2.0 delivered substantial improvements in transcript recovery and increased the number of cells available for downstream single-cell analysis. The magnitude of improvement was greatest in lower-quality samples, where sensitivity gains were most pronounced. These advances expand the range of specimens suitable for MERFISH analysis and enable broader application in clinically relevant contexts.

Importantly, enhanced sensitivity translated into deeper biological insights. MERFISH 2.0 not only increased total cell recovery but also improved detection of rare cell subtypes. In tumor samples, it enabled more comprehensive immunoprofiling, stronger inference of cell-cell interactions, and improved gene–gene correlation analysis within the tumor microenvironment. Collectively, these improvements support more detailed and biologically meaningful characterization of tissue architecture, including refined interrogation of tumor-immune and tumor-stromal organization in situ. Consistent with these observations, recent independent studies using MERFISH 2.0 in tumor and clinical tissues report substantial gain in transcript detection efficiency and improved spatial resolution of tumor-immune interactions, underscoring the expanded analytical capability of the optimized chemistry^68,69^.

Several limitations should be considered. First, performance gains are expected to vary according to tissue type, fixation protocol, specimen age, and gene panel composition. Future studies should stratify results by these pre-analytical variables and include specimen-level replication.

Second, segmentation parameters and QC thresholds influence transcript-per-cell metrics and downstream single-cell interpretations; standardized reporting of these parameters would improve cross-study comparability. Finally, this study focused primarily on top-level performance metrics between chemistries and did not pursue in-depth biological characterization. Additional differences between MERFISH 1.0 and 2.0 may emerge with deeper analysis.

Overall, MERFISH 2.0 extends the robustness of targeted spatial transcriptomics to challenging and clinically relevant samples and enables more consistent spatial gene expression profiling across diverse preservation conditions.

## Material and Methods

### Tissue samples

Fresh frozen mouse brain was collected from 6-week-old female C57BL/6 mice and snap frozen at −80°C in OCT by tissue provider BioIVT. Fixed frozen mouse brain liver, heart, kidney, lung, small intestine, spleen samples were purchased from BioIVT, and they were collected by perfusing the animal with 4% PFA first. The samples were then extracted and further fixed at 4% PFA overnight at 4°C. After sucrose gradient treatment, the samples were embedded in OCT and snap frozen at −80°C. Fresh frozen human brain tissue blocks were purchased from BioIVT and preserved at −80°C in OCT blocks. Frozen samples were cut into 10 µm thickness via a cryostat (Leica CM3050S) and fixed with 4% PFA, with mouse brain fixed at room temperature for 15 minutes, and human brain fixed at 47°C for 30 minutes. No additional fixation was performed on fixed frozen tissue slices. Afterwards, the tissue slices were washed with PBS and permeabilized with 70% ethanol at 4°C overnight before sample processing according to Vizgen’s histology guide for frozen samples (Vizgen 91600129).

FFPE human breast cancer, lung cancer from multiple patients were purchased from BioIVT and re-embedded as a tissue microarray. FFPE TMA samples were cut into 5 µm thick slices via a microtome (Leica RM2155) and placed on MERSCOPE slides according to Vizgen’s histology guide for FFPE samples (Vizgen 91600126).

### MERFISH gene panels

To evaluate the performance of MERFISH 2.0, we designed five 815-plex MERFISH 2.0 gene panels using the Vizgen’s gene pane design portal (https://portal.vizgen.com/): MERSCOPE Mouse Pan Neuro Panel 815 Gene, V 2.0 (Vizgen 10400182), MERSCOPE Human ImmunoOncology Panel 815 Gene, V 2.0 (Vizgen 10400184), MERSCOPE Human Brain Panel 815 Gene, V 2.0 (Vizgen 10400186), MERSCOPE Human Breast Cancer Panel 815 Gene, V 2.0 (Vizgen 10400188), MERSCOPE Pan Mouse Panel 815 Gene, V 2.0 (Vizgen 10400190). We used the same target regions from the MERFISH 2.0 gene panels and designed a corresponding MER-FISH 1.0 panel for each gene list. Unlike MERFISH 1.0 gene panel design, where readout bits are directly conjugated with the target region, MERFISH 2.0 encoding probes use an overhang that is conjugated with the target region and bind with the enhancers for detection. This design allows for controlled signal amplification during MERFISH 2.0 imaging. MERSCOPE Pan-Neuro Panel 815 Gene (Mouse) was used for imaging fresh frozen mouse brain samples, MER-SCOPE Pan Mouse Panel 815 Gene (Mouse) was used for imaging fixed mouse brain, liver, heart, spleen, kidney, small intestine, and lung samples. MERSCOPE Human Brain Panel 815 Gene was used for imaging fresh frozen human brain samples. MERSCOPE Breast Cancer Panel 815 Gene (Human) was used for imaging FFPE human breast cancer TMA, MERSCOPE ImmunoOncology Panel 815 Gene (Human) was used for imaging FFPE human lung cancer TMA samples.

### MERFISH 1.0 sample preparation and imaging

MERFISH 1.0 sample preparation followed published protocols and manufacturer’s instructions: MERSCOPE Fresh and Fixed Frozen Tissue Sample Preparation User Guide (Vizgen PN 91600002) and MERSCOPE FFPE Tissue Sample Preparation User Guide (Vizgen PN 91600112).

Briefly, frozen samples were stained for cell boundary using Vizgen’s Cell Boundary Kit (10400009) and later hybridized with a custom designed MERSCOPE Gene Panel Mix (Vizgen 20300008) at 37°C incubator for 36-48 hours. Following incubation, the tissues were washed with 5mL Formamide Wash Buffer at 47°C for 30 minutes, twice and embedded into a hydrogel using the Gel Embedding Premix (Vizgen 20300004), ammonium persulfate (Sigma, 09913-100G) and TEMED (N,N,N’,N’-tetramethylethylenediamine) (Sigma, T7024-25ML) from the MERSCOPE Sample Prep Kit (10400012). After the gel mix solution solidified, the samples were cleared with Clearing Solution consisting of 50uL of Proteinase K, Molecular Biology Grade (NEB, P8107S) and 5mL of Clearing Premix (Vizgen 20300003) at 37°C overnight. After removing clearing solution, the sample was washed with Sample Prep Wash Buffer and subsequently stained with DAPI and Poly T Reagent (Vizgen 20300021) for 15 minutes at room temperature. Lastly, the sample was washed for 10 minutes with 5ml of Formamide Wash Buffer and then imaged on the MERSCOPE (Vizgen 10000001) and MERSCOPE Ultra system (Vizgen 10000108).

For FFPE samples, FFPE tissue sections were dried for 20 minutes at RT and 10 minutes at 60°C. The tissue sections were then deparaffinized by Deparaffinization Buffer (Vizgen 20300112) at 55°C for 5 minutes twice, followed by another 5 minutes at RT. The deparaffinized samples were then washed with 100% ethanol three times, each 2 minutes, followed by a 2-minute 90% ethanol and a 2-minute 70% ethanol rehydration step. The rehydrated tissue sections were incubated with Decrosslinking Buffer (Vizgen 20300115) at 90°C for 15 min and cooled on bench for 5 minutes. The decrosslinked tissue sections were then incubated with Conditioning Buffer (Vizgen 20300116) at 37°C for 30 minutes, and Pre-Anchoring Reaction Buffer (Vizgen 20300113) for 2 hours at 37°C. Following anchoring pretreatment, tissue slices were stained were stained for cell boundary using Vizgen’s Cell Boundary Kit (10400009) following Vizgen’s User Guide similarly as described above (https://vizgen.com/resources/user-guides/). The sample was then washed with Formamide Wash Buffer (Vizgen 20300002) at 37°C for 30 minutes and incubated with Anchoring Buffer (Vizgen 20300117) at 37°C overnight. Following overnight incubation, the sample was washed with Formamide Wash Buffer and then gel embedded similarly to fresh frozen samples except that the samples were cleared at 47°C. After tissue clearing, the sample was treated with MERSCOPE Photobleacher (Vizgen 10100003) for 3 hours, washed with Formamide Wash Buffer at 37°C for 30 minutes, and then incubated with MERSCOPE Gene Panel Mix for 2 days at 37°C. The sample was then imaged similarly as fresh frozen samples.

### MERFISH 2.0 sample preparation and imaging

MERFISH 2.0 sample preparation followed the MERFISH 2.0 protocol from the vendor: MERSCOPE MERFISH 2.0 Sample Prep User Guide (PN 91600132). Briefly, frozen tissue sections were fixed and permeabilized via 70% ethanol overnight at 4°C, while FFPE tissue sections were deparaffinized by Deparaffinization Buffer (Vizgen 20300112) at 55°C for 5 minutes twice, followed by another 5 minutes at RT. Next, the samples were washed with 100% ethanol three times, each 2 minutes, followed by a 2-minute 90% ethanol and a 2-minute 70% ethanol rehydration step. The rehydrated tissue sections were incubated with Decrosslinking Buffer (Vizgen 20300115) at 90°C for 15 minutes and cooled on bench for 5 minutes. The permeabilized frozen or decrosslinked FFPE tissue sections were then stained for cell boundary using Vizgen’s Cell Boundary Kit (Vizgen 10400118) following Vizgen’s MERFISH 2.0 Sample Preparation User Guide for Sectioned Tissue Samples. After fixing the samples with 4% PFA for 15 minutes, the tissue sections were incubated with Conditioning Buffer (Vizgen 20300116) supplemented with RNase inhibitor at RT for 15 minutes, and then with Pre-Anchoring Reaction Buffer (Vizgen 20300113) overnight at RT in a humidified chamber. Following the overnight anchoring pretreatment, tissue slices were then briefly washed with Sample Prep Wash Buffer (Vizgen, 20300001), and Formamide Wash Buffer for 15 minutes at 37°C (Vizgen, 20300002). The samples were subsequently incubated with Anchoring Buffer (PN 20300117) at 37°C for 2 hours. Afterwards, the samples were washed with Sample Prep Wash Buffer briefly and then gel embedded and cleared using Clearing Premix supplemented with 1:100 proteinase K (NEB) overnight at 47°C. After tissue clearing, the samples were photobleached using MERSCOPE Photobleacher (Vizgen 10100003) for 3 hours, washed with Formamide Wash Buffer at 37°C for 30 minutes, and then incubated with MERSCOPE Gene Panel Mix V2.0 at 47°C overnight. The samples were then incubated with Enhancer Probe mix (Vizgen, 30300491) at 37°C overnight, and washed with Enhancer Wash Buffer (Vizgen, 20300192) at 37°C for 20 minutes, twice. The samples were stained with DAPI and Poly T Reagent V2.0 for 15 minutes at RT, washed for 10 minutes with 5ml of Formamide Wash Buffer, and then imaged on the MERSCOPE and MERSCOPE Ultra system. After image acquisition, the MERFISH data was processed by MERSCOPE and Cellpose algorithm^70^ was used to perform cell segmentation based on cell boundary staining from Vizgen’s Cell Boundary Kit. A fully detailed step-by-step instruction on the MERFISH sample prep can be found in Vizgen’s MERFISH 2.0 Sample Preparation User Guide for Sectioned Tissue Samples (Vizgen, 91600132). Full Instrumentation protocol can be found in MERSCOPE Instrument Guide (Vizgen, 9160001).

### Single cell analysis

After cell segmentation, MERSCOPE data was analyzed through single cell analysis pipeline Scanpy^71^. Areas lacking nuclear signal or exhibiting imaging artifacts were excluded from downstream analyses. For quality control filtering, cells with fewer than 10 detected transcripts were excluded from downstream analyses. We used corrected transcript counts per 100 µm² tissue area and transcript counts per cell to evaluate the assay’s sensitivity and performance. To calculate transcript density per 100 µm², the tissue area was divided into 10 × 10 µm grids. Grids containing fewer than 3 transcripts were excluded from analysis to remove low-signal regions.

The total number of transcripts within the remaining grids was summed and divided by the corresponding total area to obtain transcripts per 100 µm².

The single-cell analysis was performed with Scanpy 1.9.1. In brief, the cells from MERFISH 1.0 and MERFISH 2.0 were combined, and the cells with >= 10 transcripts were kept for clustering analysis. Harmony was used to remove potential batch effect^72^. For mouse brain, n_neighbors=15, n_pcs=50, resolution=2.0 was used; for human breast cancer tissue, n_neighbors=15, n_pcs=30, resolution=2.0 was used for leiden clustering. The resulting clusters were annotated based on their correlation with reference single-cell RNA-seq data. The spatial enrichment analysis of different cell cluster was performed with Squidpy 1.2.3^73^.

## Acknowledgements

We thank Vizgen marketing and operations team for supporting the study, and Xiaowei Zhuang and Jeffrey Moffitt for helpful discussions. This technology has been the subject of a patent application.

## Author contributions

J.H. and G.E. conceived the project. L.H, B.W., R.C., B.Y. S.G.T, L.M., T.Z., S.R., S.P., Y.S., Y.S., M.L., Y.C., L.W., N.P., M.R. performed the experiment, T.W. performed decoding analysis, S.W., H.W. designed the gene panels, R.C, H.W. performed the data analysis, A.V., M.R., J.H. wrote the manuscript.

